# Is it me or the train moving? Humans resolve sensory conflicts with a nonlinear feedback mechanism in balance control

**DOI:** 10.1101/2024.08.30.609158

**Authors:** Lorenz Assländer, Matthias Albrecht, Markus Gruber, Robert J. Peterka

## Abstract

Humans use multiple sensory systems to estimate body orientation in space. Sensory contributions change depending on context. A predominant concept for the underlying multisensory integration (MSI) is the linear summation of weighted inputs from individual sensory systems. Changes of sensory contributions are typically attributed to some mechanism explicitly adjusting weighting factors. We provide evidence for a conceptually different mechanism that performs a multisensory correction if the reference of a sensory input moves in space without the need to explicitly change sensory weights. The correction is based on a reconstruction of the sensory reference frame motion (RFM) and automatically corrects erroneous inputs, e.g., when looking at a moving train. The proposed RFM estimator contains a nonlinear dead-zone that blocks corrections at slow velocities. We first demonstrate that this mechanism accounts for the apparent changes in sensory contributions. Secondly, using a balance control model, we show predictions of specific distortions in body sway responses to perturbations caused by this nonlinearity. Experiments measuring sway responses of 24 subjects (13 female, 11 male) to visual scene movements confirmed these predictions. The findings indicate that the central nervous system resolves sensory conflicts by an internal reconstruction of the cause of the conflict. Thus, the mechanism links the concept of causal inference to shifts in sensory contributions, providing a cohesive picture of MSI for the estimation of body orientation in space.

**Significance Statement:** How the central nervous system (CNS) constructs body orientation in space from multiple sensory inputs is a fundamental question in neuroscience. It is a prerequisite to maintain balance, navigate and interact with the world. To estimate body orientation, the CNS dynamically changes the contribution of individual sensory inputs depending on context and reliability of the cues. However, it is not clear how the CNS achieves these dynamic changes. The findings in our study resolve major aspects of this question. Importantly, the proposed solution using nonlinear multisensory feedback contrasts with traditional approaches assuming context-dependent gain-scaling of individual inputs. Thus, our findings demonstrate how complex, intelligent, and unintuitive behavior can emerge from a comparably simple nonlinear feedback mechanism.

## 1 Introduction

We experience the world through multiple sensory inputs. The central nervous system (CNS) combines these inputs to create an internal representation of the world (Imamizu et al., 2000; Laurens and Angelaki, 2017; Wolpert et al., 1995). To avoid ambiguity, the CNS must resolve conflicts between individual sensory inputs. This process is called ‘causal inference’, by which changes in the external world are inferred from a given set of sensory inputs (Körding et al., 2007). Conflicting sensory inputs are a challenge for MSI in human balance control. For example, visual cues contribute to balance control. However, visual cues do not always accurately encode orientation in space. Consider standing next to a train leaving the station. In this situation, relying on vision for balance control would lead us to misinterpret the moving train as self-motion. The resulting balance corrections would induce movement with the train and lead to a fall. However, normally we do not fall. This reflects the ability of the central nervous system to modulate the contribution of visual cues used to estimate our body orientation in space and maintain balance (Horak and Macpherson, 1996; Lee and Lishman, 1975). Crucially, this selective integration involves a continuous adjustment of sensory contributions depending on context and availability — a process often attributed to sensory reweighting (Nashner and Berthoz, 1978; Peterka, 2002).

To conceptually understand the underlying MSI processes, sensory inputs can be simplified as signals encoding body orientation with respect to sensory reference frames. In free stance, the reference frames are the support surface, the visual scene, and gravito-inertial space (Peterka, 2002). Creating conflicts between inputs is a classic experimental paradigm used in MSI studies. As in the example with the moving train, sensory conflicts are also ecologically relevant as they occur in everyday life. Experimentally, RFM paradigms have been used extensively to study balance control (Pasma et al., 2015; Peterka, 2002; Kiemel et al., 2008), perception (Fetsch et al., 2009, 2012; Riecke et al., 2023) and navigation (Noel and Angelaki, 2022). These studies have shown that MSI can be represented as a weighted linear sum of individual sensory inputs. Consistent with this, CNS recordings have shown that the firing rates of multisensory neurons matched a weighted linear sum of their inputs (Cullen, 2019; Fetsch et al., 2012). However, as in the moving train example, context-dependent reweighting implies an underlying nonlinear processing underlying the change of sensory contributions (Fetsch et al., 2012; Morgan et al., 2008; Peterka, 2002).

The popular framework of optimal estimation suggests that minimizing variability in the estimate is a major criterion for adjusting sensory contributions (Dokka et al., 2010; van der Kooij and Peterka, 2011; Morgan et al., 2008; Noel and Angelaki, 2022; Ernst and Banks, 2002). However, this criterion itself does not explain how the central nervous system achieves reweighting (Angelaki et al., 2009). Furthermore, reweighting must occur rapidly in closed-loop processes (Noel et al., 2023), e.g. when a reference frame starts to move. Several models containing dynamic reweighting mechanisms for human balance control have been proposed (van der Kooij et al., 2001; Carver et al., 2006; Mahboobin et al., 2005). One of these models used a Kalman filter, which uses the minimization of noise as a criterion to select weights (van der Kooij et al., 2001). Another approach used minimization of ankle torque as a criterion (Carver et al., 2006).

In this study, we provide evidence for a closed-loop MSI mechanism that automatically adjusts sensory contributions without changing sensory weights. The mechanism was first proposed by Mergner (Mergner et al., 1991) and used as part of a wider concept for balance control (Mergner, 2010). The model proposed here started from the Mergner model and used a reductionist approach to distill the key mechanism underlying changes in sensory contributions. The mechanism fuses multiple sensory signals to estimate the movement of reference frames in space, relative to which sensory information is encoded. If the estimated movement exceeds a velocity threshold, the estimate attenuates the contribution of the respective sensory system.

We tested this mechanism for balance disturbances evoked by visual scene motion. Our results show that this mechanism is able to explain and predict sway responses of a standing human to movements of a visual reference that were previously attributed to sensory reweighting. We further rationalized that this nonlinearity must lead to specific, experimentally testable nonlinear distortions in the sway responses. In the present work we identified and experimentally confirmed such distortions providing considerable evidence for the potential existence of such a mechanism in the CNS.

## 2 Methods

### Subjects

Twenty-four subjects (13 female, 11 male, 23.3 ± 3.4 years; 172.7 ± 9.4 cm; 75.2 ± 10.9 kg) without any known balance disorders completed the study. Subjects gave written informed consent and were reimbursed with 20€for participating in the study. The protocol was approved by the University of Konstanz IRB and was in agreement with the Declaration of Helsinki.

### Experimental setup

Subjects stood upright wearing a virtual reality head-mounted display (HMD; Vive Pro Eye, HTC, Taoyuan, Taiwan) and had motion tracking markers (Vive Tracker, HTC, Taoyuan, Taiwan) attached between the shoulder blades and at greater trochanter height using Velcro straps. Subjects viewed a half-cylindrical screen (radius 1m) with high contrasting patterns (Figure 1b). The screen was tilting in an anterior-posterior direction, approximately around the ankle joint axes (8.8 cm above the floor) following predefined waveforms. Marker positions were recorded with every display update at *≈*90 Hz. Virtual environment and recording were implemented in a custom written software in Unity (Unity Technologies, San Francisco, USA) and SteamVR (Valve, Bellevue, USA) and generates balance behavior equivalent to real-world experiments (Assländer et al., 2023). The software is freely available (https://github.com/PostureControlLab/anaropia-rfm/releases). Motion smoothing and automatic resolution were enabled in SteamVR to ensure a stable framerate during the whole experiment. Pseudorandom stimuli were constructed from 80-state ternary maximum-length sequences (Davies, 1970; Peterka, 2002). The stimuli were created using a Matlab function ‘mseq’ (Buračas and Boynton, 2002) with the input options (baseVal=3, powerVal=4, shift=59). Four pseudorandom stimuli had a state duration of 0.25 s resulting in 20-s long cycles which were integrated and repeated 15 consecutive times, the fifth pseudorandom stimulus had a state duration of 0.125 s resulting in 10-s long cycles repeated 30 times (Figure 1e & f). Peak-to-peak amplitudes were 0.2°, 1°, 3°, and 5° for the 20 s cycle duration stimuli and 0.5° for the 10 s cycle duration stimulus. Two different sine stimuli were used with durations of 380 s (Figure 2); one single sine stimulus (2°, 0.2 Hz) and the sum of two sines (1°, 0.2 Hz and 1°, 0.25 Hz with +23° phase shift). The protocol included three stimuli not included in this manuscript. Following a 120 s long warm-up test, with visual screen movements to familiarize subjects with the setup, the 10 stimulus conditions were presented in random order.

**Figure 1:**
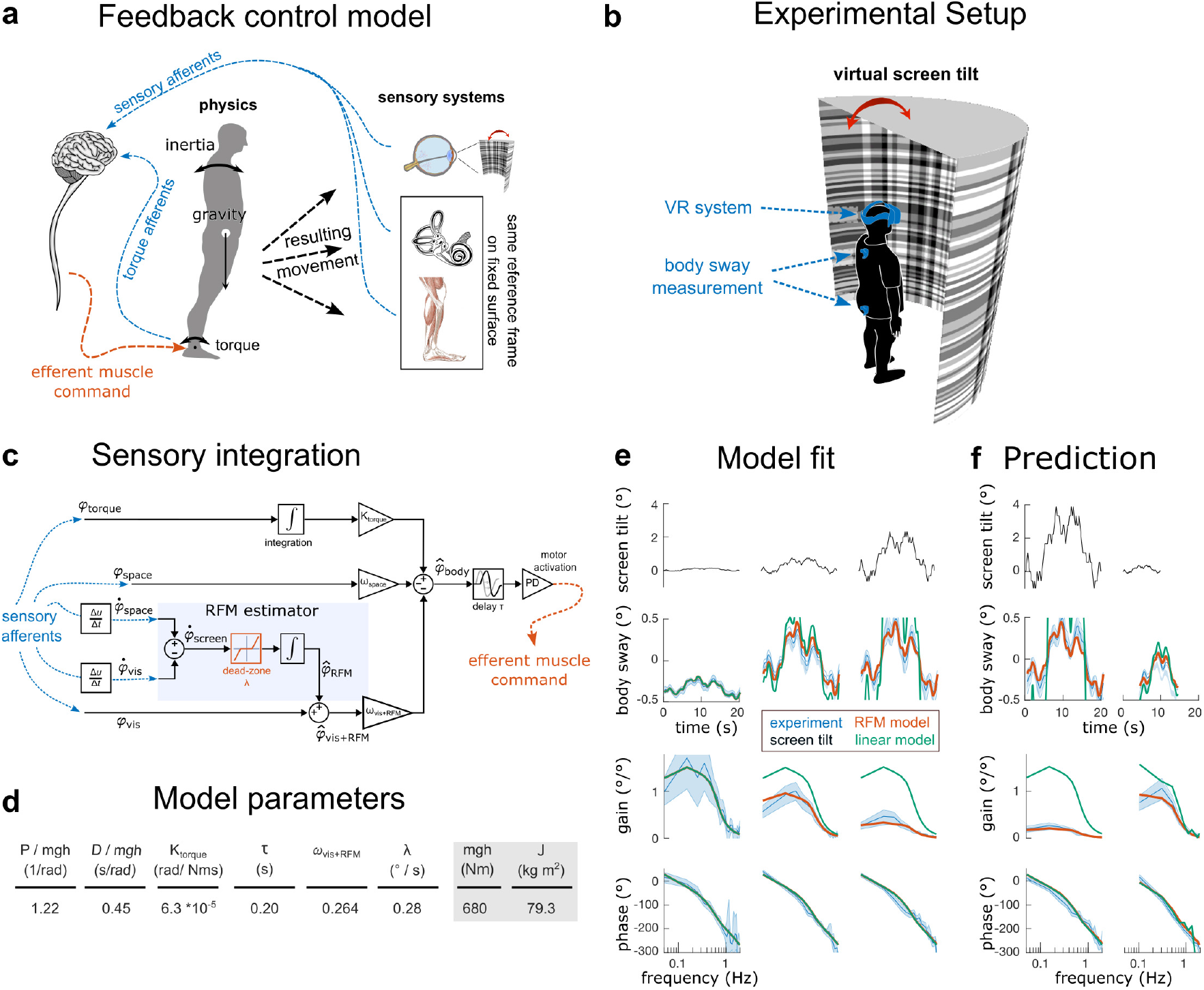
Conceptual balance control model and MSI mechanism, experimental setup model fits and model predictions. a) Conceptual balance control model. Sensory inputs encoding body orientation relative to the visual scene and relative to space, as well as ankle torque. CNS integrates sensory information and generates ankle torque, correcting deviations from the desired upright position. Corrective ankle torque accelerates the body, thereby modulating the sensory inputs, making it a closed loop system. b) Experimental setup. c) Multisensory integration mechanism. d) Model parameters estimated from experimental data: proportional (P) and derivative (D) motor activation-gains, positive torque feedback gain (K_torque_), time delay (*τ*), the visual+RFM weight (*ω*_*vis*+*RF M*_), and the dead-zone velocity threshold (*λ*; given in *°/s* for convenience). The space-reference feedback weight was derived using the convention *ω*_*vis*+*RF M*_ + *ω*_*space*_ = 1. Inverted pedulum parameters were estimated from anthropometric measures of subjects (grey shade; mass, center of mass height, and movement of inertia). e) Visual scene tilt waveform used for experimental sway, model sway predictions, and model fits. Experimental body sway averages (blue) are shown with shaded 95% bootstrap confidence bounds. Sway responses from the nonlinear model shown in c (orange) and from a linear model for comparison (green). The linear model was identical to the model in c except that the RFM estimator was removed. f) Predicted sway responses. Sway responses to stimuli that were not used for parameter fits (orange) with the experimental sway responses (blue) and linear model (green) for comparison.

**Figure 2:**
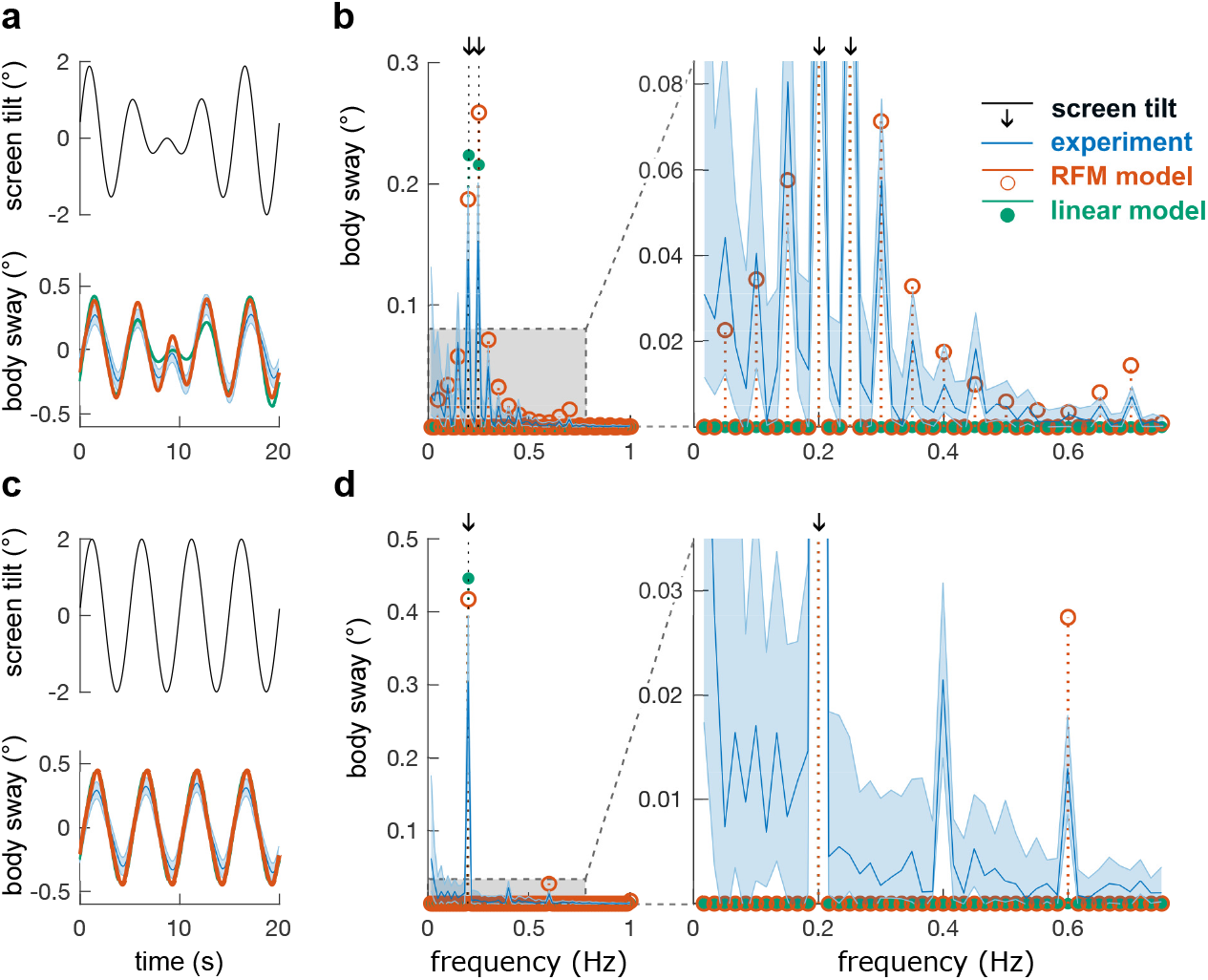
Model predictions and experimental results for sinusoidal visual scene movements. a) A waxing and waning stimulus (black) and sway responses predicted by the nonlinear model (orange) and experimental sway responses (blue shade). Linear model with the RFM mechanism removed and the linear visual weight set to 0.04 to roughly match experimental sway response amplitudes for comparison (green). b) Amplitude spectra of model predictions (orange circles) and of experimental sway responses (blue) with 95% bootstrap confidence bounds. c) Time domain and d) amplitude spectra of experimental and model-predicted (orange) sway responses to a sinusoidal 0.2 Hz stimulus. Linear model behavior is shown for comparison (green).

### Data analysis

Raw data, scripts and model used for analysis are available at the doi 10.17605/OSF.IO/EMSKV. We resampled the marker data to ensure a consistent 90 Hz sampling rate and calculated the whole-body center of mass sway using the subjects’ height, weight and anthropometric tables (Assländer et al., 2023). The first 20 s of each trial was discarded to avoid transients and data were averaged across 14 cycle repetitions and 24 subjects for each amplitude. FRFs were calculated for pseudorandom stimuli by dividing average cross-power spectra of body sway and stimulus by the average stimulus power-spectra (Bendat and Piersol, 2000). FRF calculations additionally included averaging across frequencies to obtain FRFs with approximately logarithmic frequency spacing (Peterka, 2002). For sine stimuli we averaged across the 432 single stimulus cycles to obtain across-subject time domain average responses (Figure 2, left column). Spectra were calculated from 60-s duration data segments to obtain the desired spectral frequency resolution of 1/60 s = 0.017 Hz and averaged across six repetitions per subject and 24 subjects. For time domain and spectral data, we obtained 95% confidence bounds using 1000 bootstraps of single subject average values and repeating the above described calculations. The 25th and 975th largest bootstrap values were used as upper and lower population-mean confidence bounds, respectively (Zoubir and Boashash, 1998).

### Model simulations

The model was built in Simulink 2023b (The Mathworks, Natick, USA) in two phases. In the first phase, we conducted a wide range of simulations and pilot experiments to derive the final experimental protocol. Our goal was to create a set of stimuli that allowed identification of model parameters and then use the tuned model to predict responses to additional stimuli. For the predictions we exploratorily searched for stimulus shapes, where the untuned pilot-phase model predicted strong and testable distortions in the sway responses. We found such strong distortions in the sum of two sines stimulus showing waxing and waning behavior described above, as well as a smaller distortion in a pure sine stimulus, which we then used for our final experimental protocol.

For the model presented here, we then used average subject anthropometrics to estimate moment of inertia and center of mass height. We fit the other model parameters to experimental frequency response functions *H*_*e*_ formulating the optimization problem as

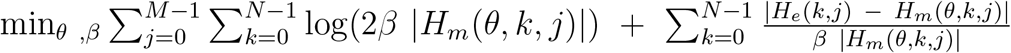

with the experimental and model FRFs *H*_*e*_ and *H*_*m*_, the N frequencies k, the M=3 pseudorandom stimulus conditions j, and the parameter vector *θ*, containing the scale parameter of the Laplace distribution *β* (Assländer et al., 2023) *and six model parameters P, D, K*_*torque*_, *τ, λ, ω*_*vis*+*RF M*_. We used a global optimization approach with the function ‘GloabalSearch’ (‘Global Optimization Toolbox’, Matlab 2023b, Mathworks, Natick, USA) and the algorithm ‘fmincon’. Parameter bounds were set to physiologically plausible values and were never reached during fitting procedures. To speed up the optimization process, we formulated the model as a transfer function to avoid using computational intensive Simulink simulations during the optimization. As a nonlinear dead-zone does not have a Laplace transform, we used an approximation developed for this purpose. The RFM estimate entering the dead-zone is an internal reconstruction of the stimulus applied during the experiment. Thus, the effect of the dead-zone can be approximated by calculating transfer functions *H*_*deadzone*_(*M, k, λ*) dividing spectra of dead-zone input 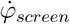 and output 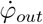 calculated from

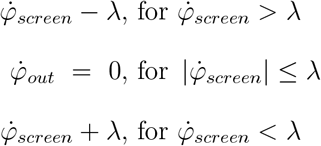

*H*_*deadzone*_(*M, k, λ*) was calculated for *λ* from 0 to 1°/s in steps of 0.01°/s. For intermediate *λ* values, we calculated the weighted average of adjacent *λ* entries in the *H*_*deadzone*_(*M, k, λ*) table. Then the model shown in Figure 1 can be formulated as a dimensionless transfer function in the frequency domain using

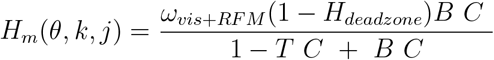

with the body dynamics 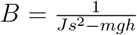 with Laplace variable s, moment of inertia J, body mass m (subtracting 2.9% for the feet), center of mass height above ankle joints h and the gravitational constant g (Winter, 2009). Motor activation *C* = (*P* + *Ds*) exp^*−s τ*^ included proportional P and derivative D components of ankle torque generation and time delay *τ*. The positive torque feedback was modeled as 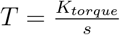. We compared the formulation of *H*_*m*_ to the Simulink model formulation to confirm their equivalence.

Following the parameter fits to the FRFs of the first three pseudorandom data sets (Figure 1e), model simulations for all five pseudorandom stimuli were run in Simulink. Then time domain and frequency domain model analyses were performed analogous to the analysis of experimental data. Linear model simulations were performed using the same procedure and the same model, fully blocking the RFM estimator path to obtain a MSI mechanism consisting of a linear weighted sum. For the prediction of nonlinear distortions, we used the model to predict sway responses to the sine and the sum of two sines stimuli with the waxing and waning behavior, again analyzing the output analogous to the experimental data. Finally, we visually extracted data from two previous studies (Mergner et al., 2005; Peterka and Benolken, 1995) to compare our model to the power law behavior described by Dokka et al. (2010). We again ran our RFM model for the sinusoidal stimuli described in these papers. As the experimental data in this step was from different subjects and obtained in different experimental setups, we manually tuned the *ω*_*vis*+*RF M*_ parameter, to 0.08 for the Mergner et al. data and 0.12 for the Peterka and Benolken data to roughly reflect the overall amplitudes of these results.

## 3 Results

### Model description

Figure 1a represents a closed loop balance control model when viewing a moving visual scene (Figure 1b). In such a situation, the central nervous system receives sensory cues from two reference frames: One sensory input encodes body orientation relative to gravito-inertial space, *φ*_*space*_, derived from vestibular and proprioceptive cues. The second sensory input encodes body orientation relative to the visual scene, *φ*_*vis*_. Additionally, the rate of change (velocity) of these two inputs, 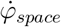 and 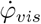, are also available. The multisensory integration mechanism (Figure 1c) combines these signals to estimate body orientation in space, 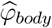, which is used to generate torque to maintain balance. However, moving the visual scene generates a conflict between the two inputs that the CNS needs to resolve to avoid falls.

To this end, we propose that one component of the MSI mechanism is an RFM estimator. The idea was derived from observations in perception and earlier studies from our laboratory (Mergner et al., 1991, 2003; Mergner, 2010; Assländer et al., 2015). Using pilot data and extensive simulations, we derived the formulation in Figure 1c as the simplest model formulation to implement this idea and to generate testable hypotheses. The model formulation was also inspired by and resembles another well established model (Peterka, 2002), to which we added the RFM estimator to account for nonlinear behavior. Within the RFM estimator, the CNS derives a signal representing the visual scene velocity in space from the two velocity inputs: 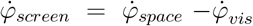. The RFM estimator processes 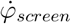 using a nonlinear dead-zone that blocks slow RFM velocities 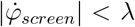 and a subsequent mathematical integration to derive the RFM estimate 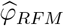. Note that 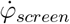 is a velocity signal, as only a velocity dead-zone in this multisensory pathway is able to account for experimental data (see below). Then 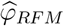 is combined with *φ*_*vis*_ to form a corrected visual contribution 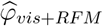. In plain words, we propose that humans use visual orientation with respect to the visual scene that is corrected by the nonlinear estimate of the visual scene movement.

A second component of the MSI mechanism forms a weighted linear sum of the corrected visual contribution 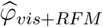 and *φ*_*space*_ with fixed weights *ω*_*vis*+*RF M*_ and *ω*_*space*_ using the convention *ω*_*vis*+*RF M*_ + *ω*_*space*_ = 1.

A third component of the MSI mechanism processes a sensory-derived measure of ankle torque *φ*_*torque*_. The ankle torque measure is mathematically integrated and scaled (*K*_*torque*_) before being summed with other sensory derived orientation information. The torque feedback is organized in a positive feedback configuration that generates a slowly acting component of ankle torque that moves the body toward an upright orientation. The torque feedback accounts for low-frequency behavior and might be considered as a self-calibration mechanism, i.e., it slowly corrects for potential errors of the dynamic feedback components (Missen et al., 2023). The torque feedback component is linear and therefore does not contribute to nonlinear changes in sensory contributions.

Considering the overall function of the MSI mechanism, a change in sensory contributions emerges when the visual scene velocity 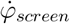 exceeds the velocity threshold *±λ* of the deadzone. Consider a simple scenario where 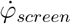 is very small 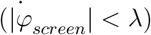. In this situation _*RF M*_ is zero and the MSI mechanism is a weighted linear sum with fixed weights. When 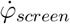 becomes larger 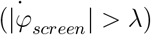, the dead-zone output is non-zero and its mathematical integral yields the RFM estimate 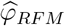, a distorted version of *φ*_*screen*_. The RFM estimate 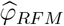 is added to the visual input *φ*_*vis*_, whenever the RFM estimate exceeds the velocity threshold *±λ*. This process effectively subtracts a nonlinearly distorted version of 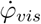 from 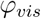 and adds a distorted version of 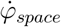. In terms of causal inference, the signal component exceeding *λ* is attributed to RFM motion by the mechanism, rather than self-motion. Thus, in this MSI mechanism, a change in sensory contributions is caused by the nonlinear RFM architecture, rather than by explicitly changing sensory weighting factors.

How much *φ*_*vis*_ is corrected depends on how much 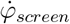 exceeds *±λ*. The system will correct in proportion to the supra-threshold component of 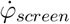. When the correction becomes very large 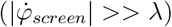, it leads to a saturation in sway responses. The reason is that 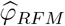 now strongly contributes, mostly canceling the visual contribution. Such saturations have been observed in several MSI studies (Mergner et al., 1991; Nashner and Berthoz, 1978; Ohshiro et al., 2011; Peterka, 2002).

### Explanatory and predictive power of the model

In a first step, we tested the ability of our model to account for sway responses across different stimulus amplitudes (Figure 1e, row 1). Sway responses to this amplitude series (Figure 1e, row 2) showed the expected less than proportional increase in sway responses compared to the increase in stimulus amplitudes (Assländer et al., 2023; Peterka, 2002). This behavior is a major nonlinear characteristic of human balance, which is typically explained by sensory reweighting. The less than proportional increase was also reflected in the frequency response functions (FRFs), which we used to characterize stimulus-response behavior in the frequency domain. The gain (sway response to stimulus amplitude ratio; Figure 1e, row 3), greatly decreased with increasing stimulus amplitudes, whereas the phase (relative timing of stimulus and response) was similar across stimulus conditions (Figure 1e, row 4).

Figure 1d shows the 6 identified model parameter values obtained using the optimization procedure. The estimated Laplace distribution width was *β* = 0.27. Notably, we fit one set of parameters for all three conditions simultaneously and parameters, including the weights *ω*_*vis*+*RF M*_, *ω*_*space*_ = 1 *− ω*_*vis*+*RF M*_, and *λ*, did not change across conditions. In other words, the model does not include any form of explicit sensory reweighting and all parameters reflect fixed properties of the MSI mechanism for balance control. The torque feedback gain (K_torque_), feedback delay (*τ*) as well as motor activation stiffness and damping parameters P and D (Figure 1d) were fully in line with estimates from previous studies (Assländer et al., 2015; Peterka, 2002).

The model accounted for the experimental FRF shapes at all three stimulus amplitudes (Figure 1e, rows 3 and 4) and reproduced time-domain sway responses (Figure 1e, row 2) with a variance accounted for (VAF) of 89%, 82% and 95%, respectively. For comparison, the behavior of a linear model with fixed weights and without any form of reweighting is shown in green (Figure 1e). The FRFs of the linear model do not change across conditions, reflecting the linear increase of sway amplitudes with stimulus amplitudes, leading to sway responses largely exceeding the experimental and RFM model results at larger stimulus amplitudes. As the RFM model reproduced the less than proportional change in sway response amplitudes, the result demonstrates the ability of the model to reproduce behavior typically attributed to changes of sensory weighting factors.

Predicted sway responses to the two pseudorandom visual stimuli that were not used in the original parameter identification procedure are shown in Figure 1f. Experimental results were in excellent agreement with RFM model predictions, in the frequency domain (Figure 1f, rows 3 and 4), as well as in the time domain with VAFs of 91% and 81%, respectively (Figure 1f, row 2).

### Distortions in sway responses to sinusoidal stimuli

The ability to explain and predict responses to different stimulus amplitudes constitutes considerable evidence for the validity of the proposed MSI mechanism. A second test of our model was the specific distortions we had predicted a priori to our experiments using model simulations (see Methods). The stimuli were created based on the rationale that distortions are expected in stimuli with varying velocities, as the dead-zone blocks the RFM estimate at slow velocities, but contributes at faster velocities. Therefore, the feedback dynamics would change nonlinearly within one stimulus cycle.

As sine waves continuously change velocities, we predicted that balance responses should show some form of nonlinear distortion. Our a priori simulations showed that we can maximize the effect, when using a waxing and waning stimulus pattern constructed by superimposing two closely adjacent sinusoidal frequencies (Figure 2a). For the results presented here, we used the model parameters derived from responses to the pseudorandom stimuli (Figure 1d). The predictions showed sway responses at the two stimulus frequencies (black arrows in Figure 2b), as would be expected in any linear model (green dots in Figure 2b). However, the RFM model also predicted sway responses at a number of specific frequencies other than those of the stimulus (Figure 2b; red circles with values greater than zero), representing the nonlinear distortion in the sway pattern. The experimental results were largely consistent with the predicted sway responses with spectral peaks from the experimental data occurring at the same frequencies as the model-predicted spectral peaks (Figure 2b). In the time domain, sway responses followed the predicted shape with slightly smaller experimental sway responses as predicted and a VAF of 81%. Furthermore, the experimental spectrum exhibited variable low frequency components that declined gradually with increasing frequency, aligning with spectral patterns observed during quiet stance (Laboissière et al., 2015).

We also tested whether sway responses would be distorted when using a single sine stimulus (Figure 2c). The dead-zone would again block the RFM estimate, during slow phases of the sine, thereby distorting the feedback. The predicted distortion was surprisingly small, but visible in the spectrum at 0.6 Hz (Figure 2d). In agreement with this prediction, the experimental spectrum also showed a frequency peak at 0.6 Hz. In addition, experiments showed a second small peak at 0.4 Hz, which was not predicted by the model and could be related to the simplifications used in our approach (see Discussion). Overall, the time domain sway response to the sinusoidal stimulus was predicted with a VAF of 86%.

### Power-law velocity dependence

Two earlier studies (Peterka and Benolken, 1995; Mergner et al., 2005) characterized sway responses evoked by sinusoidal visual stimuli over a range of frequencies and amplitudes, providing additional data to test the explanatory power of the RFM model. Additionally, a subset of these data from 0.2 Hz tests with stimulus velocities ranging from about 0.25 to 12.5°/s were shown to be consistent with a Bayesian attribution model (Dokka et al., 2010), also shown in Figure 3. Specifically, the Bayesian analysis reproduced a power-law relationship between response gain (ratio of sway velocity to stimulus velocity) and peak visual stimulus velocity. Since Bayesian inference is widely considered to be an organizing principle in sensorimotor processing and control, we considered it relevant to determine the extent to which the RFM model reproduces this power-law relationship.

**Figure 3:**
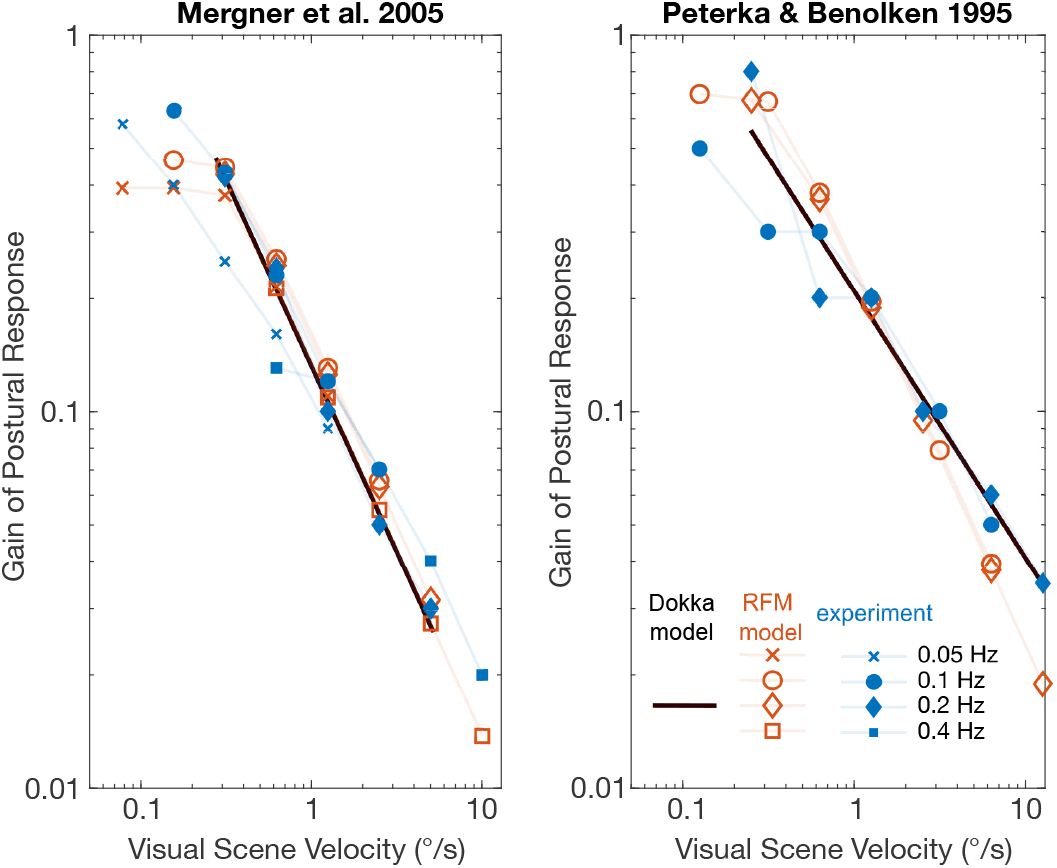
RFM model behavior, model behavior of Dokka et al. (2010) and experimental results for sinusoidal visual scene movements across multiple amplitudes and frequencies. Experimental data were manually extracted from Figure 3 in Mergner et al. (2005) and Figure 4 in Peterka and Benolken (1995) and displayed as described in Dokka et al. (2010). RFM model results are for parameters as shown in Figure 1d with the exception of the visual+RFM weight, which was set to *ω*_*vis*+*RF M*_ = 0.08 and to *ω*_*vis*+*RF M*_ = 0.12 for comparison with the Mergner et al. and Peterka and Benolken data, respectively.

Response gain and peak visual stimulus velocity data from the earlier studies showed an approximately linear relation for 0.2 Hz stimulus frequencies when plotted on log-log axes (Figure 3), as expected for power-law behavior (Dokka et al., 2010). Similar power law behavior was exhibited also for data from 0.05, 0.1, and 0.4 Hz stimulus frequencies (not included in the Dokka et al. (2010) analysis). We found that the RFM model with our previously identified parameters (Figure 1d) displayed similar power-law behavior over a wide range of stimulus amplitudes thus demonstrating behavior that has been linked to Bayesian inference.

However, the response gains from the RFM model were systematically larger than the experimental data when using the parameters estimated for our subjects and conditions (Figure 1d). By lowering the model’s visual+RFM weight, *ω*_*vis*+*RF M*_, to 0.08 and 0.12 for the Mergner et al. (2005) data and the Peterka and Benolken (1995) data, respectively, good agreement was obtained to the experimental data (Figure 3). Notably, at very low stimulus velocities below about 0.2°/s the RFM gains were lower than expected from power-law behavior.

## 4 Discussion

In the present study, we provide evidence that changes in sensory contributions to balance control emerge from a multisensory reconstruction of visual scene motion in the central nervous system. This mechanism is able to explain and predict less than proportional increases in sway response amplitudes with increasing stimulus amplitudes. The model further predicted spectral distortions with sway responses evoked at frequencies not present in the stimulus. Overall, the model was in excellent agreement, explaining and predicting experimental sway responses to a total of 7 stimuli with a VAF between 81% and 95%. In addition, the model is comparatively simple, with only 6 model parameters and a physiologically plausible architecture. Finally, we showed that the model reproduces power-law behavior across a wide range of stimulus velocities that has been related to sensory processing based on Bayesian inference principles (Dokka et al., 2010). In the following, we discuss the generality of RFM estimation and its potential role in noise reduction and causal inference.

Nonlinear RFM estimation may also play a role in other tasks that require an internal reconstruction of body orientation in space. In the current study, we used visual scene motion to induce sensory conflicts. However, the RFM estimator was inspired by models incorporating RFM-like mechanisms from previous studies. The models reproduced reweighting-like balance behavior induced by other RFMs, such as support surface tilts (Assländer et al., 2015; Mergner et al., 2003) or movements of a haptic reference (Assländer et al., 2018). The RFM concept is also consistent with human perception during sensory conflicts (Mergner et al., 1991).

Mergner et al. (Mergner et al., 1991) found that humans perceive a slow rotation of the body relative to the head as a head rotation in space, even when the body is rotating underneath the fixed head. This illusion fades with increasing velocities and the body is increasingly correctly perceived as moving in space. This perceptual pattern was accounted for by a multisensory nonlinear RFM estimation, similar to the one presented here. The concept of RFM estimation is also consistent with the train illusion in which a train that begins to move is interpreted as self-motion rather than train motion. The self-motion illusion fades as the train speeds up and the RFM estimate starts to correct the visual input.

All of these results share a common framework. For the estimation of body orientation in space the CNS

1. calculates multisensory RFM estimates,
2. uses the RFM estimates to correct sensory inputs if they exceed a certain velocity, and
3. linearly combines the corrected sensory inputs with fixed weights.

The mechanism acts rapidly, thus meeting the requirements for MSI of continuous sensory signals in closed loop control tasks (Noel et al., 2023). Thus, the evidence from studies in balance control and perception indicate that nonlinear RFM estimation could be a general mechanism to resolve sensory conflicts in MSI.

In our model, the RFM estimate adjusts sensory contributions to balance without explicitly changing sensory weighting factors. In principle, different models could be formulated that process sensory cues to produce stimulus-dependent weight adjustments. By definition these would be non-linear models that would also show non-linear stimulus-response behavior. Rather than investigating a wide variety of models, our approach was guided by the principle of Occam’s razor, thus the search for a very simple model with the potential to modulate sensory contributions. The ability of our model to account for a variety of new experimental results as well as stimulus-response behavior previously considered to be associated with Bayesian processing of sensory cues provide strong evidence that the RFM mechanism may provide a realistic representation of CNS mechanisms that process multisensory cues. Nevertheless, there is a need to consider other potential model formulations and to add components to our model, such as muscle models, that include additional nonlinearities, and to use the models to identify future experimental tests that give further insight into the validity of any model formulation.

Several aspects make the model physiologically plausible. First, its architecture only contains summations, derivatives/integrations, a time delay, gain scaling, and a dead-zone. All of these are operations and properties well known to exist on the level of individual or an assembly of neurons. Second, the model parameters (Figure 1d) provide meaningful quantities which are related behavioral and physiological aspects of sensory integration. The fixed visual+RFM and space weighting factors can be interpreted as the visual and proprioceptive/vestibular contributions to balance during unperturbed stance when sway velocities are low and remain below the nonlinear threshold. In our sample, the average contribution of visual cues was 26.7%, while vestibular+proprioceptive cues contributed 73.3%. The threshold *λ* of the nonlinear dead-zone indicates that subjects start to attenuate the visual contribution if the visual reference frame moves faster than *λ* = *±*0.28^*°*^*/s*. The model also explicitly accounts for time delays in the system and reproduces the compliant behavior characteristic for human posture control (relatively small motor activation gain). In summary, all model components are physiologically plausible and model parameters have a physiologically and behaviorally relevant meaning.

Vestibular information likely plays a dominant role in RFM estimation. In our study, subjects stood on a fixed level surface. Thus, both proprioceptive and vestibular systems provided inputs from space-veridical reference frames. Assuming that both sensory inputs are used for the RFM estimation, we should see similar reweighting effects in subjects with normal sensory function and those with bilaterally absent vestibular function. However, previous studies showed that subjects without vestibular function were essentially unable to reweight visual information when standing on a fixed support surface (Peterka, 2002; Peterka and Benolken, 1995). Re-interpreting these results with respect to the RFM mechanism, vestibular loss subjects seem to be unable to use proprioceptive cues for RFM estimations for visual scene movements. They appear to effectively use fixed sensory weights across all visual stimulus amplitudes. Thus, the gravito-inertial space reference provided by the vestibular system appears to be of primary importance for the RFM estimation.

One benefit of the RFM concept is the creation of context. For intelligent behavior, it is widely understood that the CNS utilizes an internal representation of the outside world. The RFM estimate constitutes such an internal representation of the external environment derived from the nonlinear processing of available sensory cues. Furthermore, intelligent behavior, in the sense of a context-dependent use of sensory information, emerges from this process. The underlying nonlinear processing of multi-sensory information infers changes in environmental conditions and generates rapid MSI changes that adjust for them. The RFM estimation as it applies to balance control can therefore be considered to represent the embodiment of a causal inference mechanism specifically designed to implement rapid compensation for a wide variety of sensory conflicts.

Another major benefit of combining multiple sensory inputs is the reduction of variability in estimates (Angelaki et al., 2009; Ernst and Banks, 2002; Faisal et al., 2008; Wolpert, 2007). This raises the question of whether the RFM estimator can have a noise reducing effect. While it is inherently difficult to directly assess noise properties of a sensory system, there are studies that support the notion that vestibular signals contain a high noise level (van der Kooij and Peterka, 2011; Nouri and Karmali, 2018). An earlier study showed that a velocity dead-zone can be used to partially block vestibular noise (Mergner et al., 2009). Applied to the RFM estimator in our model when the visual scene is not moving, the dead-zone would block or partially block proprioceptive/vestibular noise in the 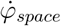 signal as well as noise in the 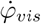 signal as the random fluctuations would be largely below the dead-zone threshold and thus not contribute to the 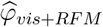 signal. In situations of a moving visual scene, however, the 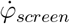 signal would cross the dead-zone threshold and additional proprioceptive/vestibular noise as well as visual noise would enter the system. The dead-zone in the RFM estimate could therefore achieve noise reduction by partially blocking noisy vestibular signals in situations of a stationary reference frame. The RFM concept would then represent a compromise between reducing noise and avoiding misinterpretations of sensory information when reference frames are moving and therefore unreliable. Such a trade-off between precision and accuracy is also captured by a Bayesian attribution model, which has been used to interpret the data presented in Figure 3 (Dokka et al., 2010). Unfortunately, a direct comparison of the RFM model and the Bayesian attribution model is not possible as the model formulations have considerable conceptual differences. Dokka et al. (2010) derived the power-law relation between gain values and stimulus velocity based on assumed noise properties and assumed an ideal transformation between sensed body motion and actual body motion. However, sensed body motion and actual body motion are linked by the dynamics of the control mechanism, which determine both gain and phase behavior. The Bayesian attribution model includes probabilistic considerations but excludes explicit control mechanisms. For example, gain and phase characteristics such as in Figure 1e and f or frequency distortions such as in Figure 2 cannot be explained without accounting for the dynamic properties of the system. In simple words, the RFM model is a control model, while the Bayesian attribution model is not. On the other hand the Bayesian model is based on stochastic considerations, while the RFM model does not take stochastic properties into account. Implementing stochastic considerations into control models is a non-trivial task that needs to be addressed in future work.

Stochastic properties might be important to consider, especially in the context of the nonlinear RFM mechanism. Sensory signals, neural processing and muscle contractions contain noise, leading to variability in body orientation estimates and resulting in random sway. This random sway component is superimposed on the sway evoked by the stimulus. Using a large amount of averaging, we reduced the random sway component in our experimental data and extracted the sway responses to the stimuli. For linear systems, handling noise in this way is a valid approach (Pintelon and Schoukens, 2012) and has been widely used in MSI research. However, for nonlinear systems, the approach could result in systematic biases. A noteworthy interaction between noise and nonlinear dead-zone elements is stochastic resonance (Benzi et al., 1981). In the context of our model, the internal estimate 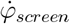 will fluctuate around the true value due to sensory noise. These fluctuations will occasionally cross the dead-zone threshold, even if 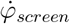 is, on average, below the threshold. Thus, sensory noise and stimulus characteristics are likely to interact unintuitively with the nonlinear dead-zone in the RFM mechanism. Explicitly accounting for internal noise sources could therefore further improve model validity and predictions. Furthermore, these interactions could be used to identify noise properties at different locations in the feedback mechanism. Including stochastic components in a nonlinear feedback-control model requires mathematical methods tailored to the problem. We used a set of established simplifications to formulate our model. These include the concept of reference frames instead of individual peripheral sensory organ signals, the concept of body schema (i.e., ‘visual’ cues encode body orientation, rather than head orientation relative to the visual scene), modeling the body as a single inverted pendulum (neglecting coupling forces and potential hip and neck movements for head stabilization; De Nunzio et al. 2005), assuming ideal transformations between position, velocity and acceleration signals, the assumption of a single lumped time delay, and noise-free sensory and central signals. Our goal was to uncover fundamental properties before adding complexity, which can make interpretations difficult if fundamental properties are not understood. For this purpose, we also constrained the range of stimulus amplitudes to avoid transient or voluntary movements, such as compensatory stepping caused by pushing subjects to their stability limits. Despite these simplifications, our model accounted for important features present in the experimental data.

Our results showed that we can create quantitative models with considerable explanatory and predictive power. Such wide experimental characterizations should also allow us to compare competing models for balance control in future studies. This includes existing models (Mergner, 2010; van der Kooij et al., 2001; Carver et al., 2006; Mahboobin et al., 2005), as well as the consideration of additional model details, such as nonlinear muscle properties (Loram et al., 2007) or muscle activation dynamics (Pasma et al., 2015). Further investigations could also determine if RFM models can predict the time course of dynamic adaptations that occur with sudden changes in stimulus conditions (Carver et al., 2006; Assländer and Peterka, 2014) and test the identified deviation from the power-law relation presented in Figure 3.

In conclusion, we have shown how a velocity dead-zone in the multisensory reconstruction of reference frame movements leads to context-dependent changes of sensory contributions to balance and predicted specific distortions in sway behavior. In addition, this mechanism reproduced a power-law stimulus-response relationship that has been related to Bayesian sensory integration principles. The mechanism can be considered to be a rapid-acting causal-inference mechanism suitable for use in a closed-loop feedback control system.

## References

Angelaki DE, Gu Y, DeAngelis GC (2009) Multisensory integration: psychophysics, neurophysiology, and computation. Current Opinion in Neurobiology 19:452–458.

Assländer L, Albrecht M, Diehl M, Missen KJ, Carpenter MG, Streuber S (2023) Estimation of the visual contribution to standing balance using virtual reality. Scientific Reports 13:2594.

Assländer L, Hettich G, Mergner T (2015) Visual contribution to human standing balance during support surface tilts. Human Movement Science 41:147–164.

Assländer L, Peterka RJ (2014) Sensory reweighting dynamics in human postural control. Journal of neurophysiology 111:1852–64.

Assländer L, Smith CP, Reynolds RF (2018) Sensory integration of a light touch reference in human standing balance. PloS one 13:e0197316–e0197316.

Bendat J, Piersol A (2000) Random Data: Analysis and Measurement Procedures, 3rd edition Wiley Series in Probability and Statistics. Wiley.

Benzi R, Sutera A, Vulpiani A (1981) The mechanism of stochastic resonance. Journal of Physics A: mathematical and general 14: L453.

Buračas GT, Boynton GM (2002) Efficient design of event-related fmri experiments using m-sequences. Neuroimage 16:801–813.

Carver S, Kiemel T, Jeka JJ (2006) Modeling the dynamics of sensory reweighting. Biological cybernetics 95:123–134.

Cullen KE (2019) Vestibular processing during natural self-motion: implications for perception and action. Nature Reviews Neuroscience 20:346–363 Publisher: Springer US.

Davies W (1970) System Identification for Self-Adaptive Control Wiley-Interscience.

De Nunzio AM, Nardone A, Schieppati M (2005) Head stabilization on a continuously oscillating platform: The effect of a proprioceptive disturbance on the balancing strategy. Experimental Brain Research 165:261–272 ISBN: 0014-4819 (Print)\n0014-4819 (Linking).

Dokka K, Kenyon RV, Keshner Ea, Kording KP (2010) Self versus environment motion in postural control. PLoS computational biology 6:e1000680–e1000680.

Ernst MO, Banks MS (2002) Humans integrate visual and haptic information in a statistically optimal fashion. Nature 415:429–433.

Faisal AA, Selen LPJ, Wolpert DM (2008) Noise in the nervous system. Nature Reviews Neuroscience 9:292–303.

Fetsch CR, Pouget A, Deangelis GC, Angelaki DE (2012) Neural correlates of reliability-based cue weighting during multisensory integration. Nature Neuroscience 15:146–154.

Fetsch CR, Turner AH, DeAngelis GC, Angelaki DE (2009) Dynamic reweighting of visual and vestibular cues during self-motion perception. The Journal of Neuroscience 29:15601–15612.

Horak F, Macpherson J (1996) Postural orientation and equilibrium In Geiger SR, editor, Handbook of Physiology - Section 12, pp. 255–292. American Physiological Soc.

Imamizu H, Miyauchi S, Tamada T, Sasaki Y, Takino R, PuÉtz B, Yoshioka T, Kawato M (2000) Human cerebellar activity reflecting an acquired internal model of a new tool. Nature 403:192–195.

Kiemel T, Elahi AJ, Jeka JJ (2008) Identification of the plant for upright stance in humans: multiple movement patterns from a single neural strategy. Journal of neurophysiology 100:3394–3406.

Körding KP, Beierholm U, Ma WJ, Quartz S, Tenenbaum JB, Shams L (2007) Causal inference in multisensory perception. PLoS ONE 2.

Laboissière R, Letievant JC, Ionescu E, Barraud PA, Mazzuca M, Cian C (2015) Relationship between spectral characteristics of spontaneous postural sway and motion sickness susceptibility. PLOS ONE 10:e0144466 Publisher: Public Library of Science.

Laurens J, Angelaki DE (2017) A unified internal model theory to resolve the paradox of active versus passive self-motion sensation. eLife 6:e28074 Publisher: eLife Sciences Publications, Ltd.

Lee DN, Lishman JR (1975) Visual proprioceptive control of stance. Journal of Human Movement Studies 1:87–95 Publisher: Teviot Scientific Publications.

Loram ID, Maganaris CN, Lakie M (2007) The passive, human calf muscles in relation to standing: the non-linear decrease from short range to long range stiffness. The Journal of physiology 584:661–675.

Mahboobin A, Loughlin PJ, Redfern MS, Sparto PJ (2005) Sensory re-weighting in human postural control during moving-scene perturbations. Experimental Brain Research 167:260–267.

Mergner T (2010) A neurological view on reactive human stance control. Annual Reviews in Control 34:177–198 Publisher: International Federation of Automatic Control.

Mergner T, Maurer C, Peterka RJ (2003) A multisensory posture control model of human upright stance. Progress in brain research 142:189–201.

Mergner T, Schweigart G, Maurer C, Blümle A (2005) Human postural responses to motion of real and virtual visual environments under different support base conditions. Experimental brain research 167:535–556.

Mergner T, Siebold C, Schweigart G, Becker W (1991) Human perception of horizontal trunk and head rotation in space during vestibular and neck stimulation. Experimental brain research 85:389–404 ISBN: 0014-4819 (Print)\n0014-4819 (Linking).

Mergner T, Schweigart G, Fennell L, Maurer C (2009) Posture control in vestibular-loss patients. Annals of the New York Academy of Sciences 1164:206–15.

Missen KJ, Assländer L, Babichuk A, Chua R, Inglis JT, Carpenter MG (2023) The role of torque feedback in standing balance. Journal of Neurophysiology 130:585–595.

Morgan ML, DeAngelis GC, Angelaki DE (2008) Multisensory integration in macaque visual cortex depends on cue reliability. Neuron 59:662–673.

Nashner L, Berthoz A (1978) Visual contribution to rapid motor responses during postural control. Brain research 150:403–7.

Noel JP, Angelaki DE (2022) Cognitive, systems, and computational neurosciences of the self in motion. Annual review of psychology 73:103–129.

Noel JP, Bill J, Ding H, Vastola J, Deangelis GC, Angelaki DE, Drugowitsch J (2023) Causal inference during closed-loop navigation: Parsing of self- and object-motion. Philosophical Transactions of the Royal Society B: Biological Sciences 378 Publisher: Royal Society Publishing.

Nouri S, Karmali F (2018) Variability in the vestibulo-ocular reflex and vestibular perception. Neuroscience 393:350–365.

Ohshiro T, Angelaki DE, Deangelis GC (2011) A normalization model of multisensory integration. Nature Neuroscience 14:775–782.

Pasma JH, Engelhart D, Maier AB, Schouten AC, van der Kooij H, Meskers CGM (2015) Changes in sensory reweighting of proprioceptive information during standing balance with age and disease. Journal of neurophysiology 114:3220–33.

Peterka RJ, Benolken MS (1995) Role of somatosensory and vestibular cues in attenuating visually induced human postural sway. Experimental brain research 105:101–110.

Peterka RJ (2002) Sensorimotor integration in human postural control. Journal of neurophysiology 88:1097–118.

Pintelon R, Schoukens J (2012) System Identification: A Frequency Domain Approach John Wiley & Sons Pages: 743.

Riecke B, Murovec B, Campos J, Keshavarz B (2023) Beyond the eye: Multisensory contributions to the sensation of illusory self-motion (vection). Multisensory Research 36:1–38.

van der Kooij H, Jacobs R, Koopman B, van der Helm F (2001) An adaptive model of sensory integration in a dynamic environment applied to human stance control. Biological cybernetics 84:103–15.

van der Kooij H, Peterka RJ (2011) Non-linear stimulus-response behavior of the human stance control system is predicted by optimization of a system with sensory and motor noise. Journal of computational neuroscience 30:759–78 ISBN: 1573-6873 (Electronic)\n0929-5313 (Linking).

Winter D (2009) Biomechanics and Motor Control of Human Movement Wiley.

Wolpert DM (2007) Probabilistic models in human sensorimotor control. Human movement science 26:511–524.

Wolpert DM, Ghahramani Z, Jordan MI (1995) An internal model for sensorimotor integration. Science 269:1880–1882.

Zoubir A, Boashash B (1998) The bootstrap and its application in signal processing. IEEE Signal Processing Magazine 15:56–76.

